# In-depth investigation of the mechanisms of high and low residual feed intake regulating hens during the late laying period via liver and gut microbiota

**DOI:** 10.1101/2024.03.20.585923

**Authors:** Zhouyang Gao, Chuanwei Zheng, Zhiqiong Mao, Jiangxia Zheng, Dan Liu, Guiyun Xu

**Affiliations:** College of Animal Science and Technology, China Agricultural University, Beijing 100193, China; Beinongda Technology Co., Ltd., Beijing 100083, China

**Keywords:** Residual feed intake, gut microbiota, livestock’s efficiency, laying hens, poultry production

## Abstract

Residual feed intake (RFI) is a more accurate indicator of feed efficiency than the feed conversion ratio (FCR) and is widely used to measure the efficiency of livestock and poultry feed utilization. Typically, Low RFI (LRFI) implies higher feed conversion efficiency, while high RFI (HRFI) indicates lower feed conversion efficiency. This study systematically explored the differences between high and low RFI and the function of the liver and cecum microbes of hens during the late laying period by multiple-omics techniques and further explored the interaction among microorganisms, the function of tissues and organs, and body metabolism. The results showed that the length and mass of the digestive organs in the LRFI group were higher than those in the HRFI group as well as the chest width. Additionally, the key genes and metabolites regulating RFI in hens during the late laying phase were found to be *ADCY2, ADCY8, CCKAR, ACSS2, FABP1, FABP4*, and LysoPI (18:2(9Z,12Z)/0:0) in the liver. The levels of AST, HDL-C and ACTH in the serum were considered candidate markers influencing RFI. By conducting a microbiome-metabolome association analysis, we have identified the dominant and beneficial microbial community in the gut of LRFI groups, such as *Oscillospirales*, *Ruminococcaceae*, and *Butyricicoccaceae*, which offers a theoretical basis for understanding how the gut microbiota regulates RFI. These results will provide a scientific basis for the molecular mechanism of RFI phenotypic variation in late laying hens.

## 1. Introduction

With the rapid development of large scale and intensive breeding modes, feed cost account for more than 70% of the total cost in the poultry production process^[1]^. The optimization of feed efficiency is one of the main challenges of livestock and poultry genetic improvement programs. Feed intake is one of the most important factors limiting production efficiency and is closely related to animal health and product quality^[2]^. Therefore, in poultry production, breeding selection with high feed efficiency can effectively reduce animal feed consumption and the total cost of production, alleviating the pressure of raw material competition and contributing to the sustainable development of environmental protection.

At present, the main evaluation of feed utilization refers to the FCR and RFI. FCR is defined as the ratio of the weight of feed intake to the weight of livestock or poultry products produced in a certain measurement period (Hess et al., 1941), which is called the feed-to-egg ratio in laying hens^[3]^. However, the researchers found that there is a nonlinear relationship between FCR and its components and with complex additional factors and multiplicative effects^[4]^, so direct selection of FCR may not be the best way to breed improvement. RFI^[5]^ (Koch et al.,1963) is defined as the difference between the actual feed intake of livestock or poultry and the expected feed intake for maintenance and growth needs, which is considered to be a more suitable indicator for measuring the feed utilization efficiency of livestock and poultry. A relevant study found that the selection of hens with lower RFI as the feed utilization index can significantly reduce feed intake without changing the weight of the group, thereby effectively saving production costs^[6]^.

Poultry feeding behavior is a large, complex and elaborate neural network that includes various classes of hormones, neurotransmitters, cytokines, and secreted factors^[7]^. The feeding behavior of poultry is inseparable from its unique physiological structure and digestive organ composition. The regulation of poultry feeding behavior is mainly composed of the central nervous system, peripheral organs and feeding behavior is stopped by negative feedback regulation. The central nervous system is mainly controlled by the hypothalamus of poultry, while peripheral organs include sensory organs, digestive tract, liver and adipose tissue, which are linked to the central nervous system through the vagus nerve, hormones, nutrients or metabolites in the blood and regulate appetite by secreting various peptides and steroid hormones^[8, 9]^.

The digestive system of poultry mainly includes the craw, proventriculus, gizzard, small intestine (including the duodenum, jejunum, and ileum), large intestine (including the cecum and rectum) and cloaca^[10]^. The glandular stomach and muscle stomach are considered to be the “true stomach” of chickens, the glandular stomach is the part of the digestive tract that secretes pepsin and hydrochloric acid and is mainly responsible for chemical digestion, and the muscle stomach is mainly responsible for mechanical grinding and squeezing chyme^[11]^. The small intestine is the main place for the digestion and absorption of nutrients, and the efficiency of digestion and absorption of feed nutrients is increased through rapid peristalsis. A study has shown that chickens with LRFI have longer duodenal lengths and are more digestible and absorbed^[12]^. The cecum is the main site for microbial fermentation to degrade indigestible nutrients such as crude fiber, absorbing electrolytes and water^[13]^. The study found that through 16S sequencing, RFI was closely related to the composition of intestinal microorganisms, and the abundance of beneficial flora in LRFI chickens was significantly improved^[14]^. The liver is the largest substantial organ in poultry and is responsible for a range of physiological and biochemical functions, such as sugar metabolism, lipid metabolism, protein synthesis and metabolism^[15]^. It has been reported that transcriptome sequencing of chicken livers with different feed efficiency has found that the differential genes with high and low feed utilization is related to appetite, cell activity and lipid metabolism. The utilization of energy is mainly optimized by reducing cell activity, immune response and energy expenditure^[16]^. In addition, systematic concentrations of key blood parameters related to feeding, growth, nutrient distribution and utilization have been found to be potential physiological markers of RFI, mainly including biochemical metabolites and various circulating hormones. There are many studies on the correlation between RFI and blood parameters in livestock and poultry^[17–19]^.

China is the world’s largest producer and consumer of chicken eggs, and the current goal of the industry is to extend the production cycle of laying hens to 100 weeks of age to improve the sustainability of poultry production. Currently, there is tremendous potential for the application of multiomics technologies in livestock production. Through multiomics technologies, we can gain an in-depth understanding of the genetic characteristics and nutritional requirements of poultry. However, the mechanism of RFI in laying hens during the late period remains unclear. Above all, this study mainly measured the serum physiological and biochemical indices, digestive organ parameters, and multiomics technology of hens with HRFI and LRFI in the late stage of hens and further explored the interaction between microorganisms, tissues, organ functions and body metabolism, which provided a certain theoretical basis for RFI selection in laying hens during the late stage.

## 2. Materials and methods

### 2.1 Animal experimental design

The entire trial period was 27 to 70 weeks. A total of 1557 hens from a pedigreed line of Rhode Island Red were used from Beinongda Technology Co., Ltd., China. In the trial, all chickens were in the same environment and kept in separate cages, with the same feed and drinking water freely. A 16-hour light and 8-hour dark cycle (16L:8D) was performed. According to the feeding standards of NRC^[20]^, corn and soybean meal were selected as the main dietary components and were formed according to the appropriate proportions in Table 1. All hens remained in good health during the whole feeding period.

**Table 1.**
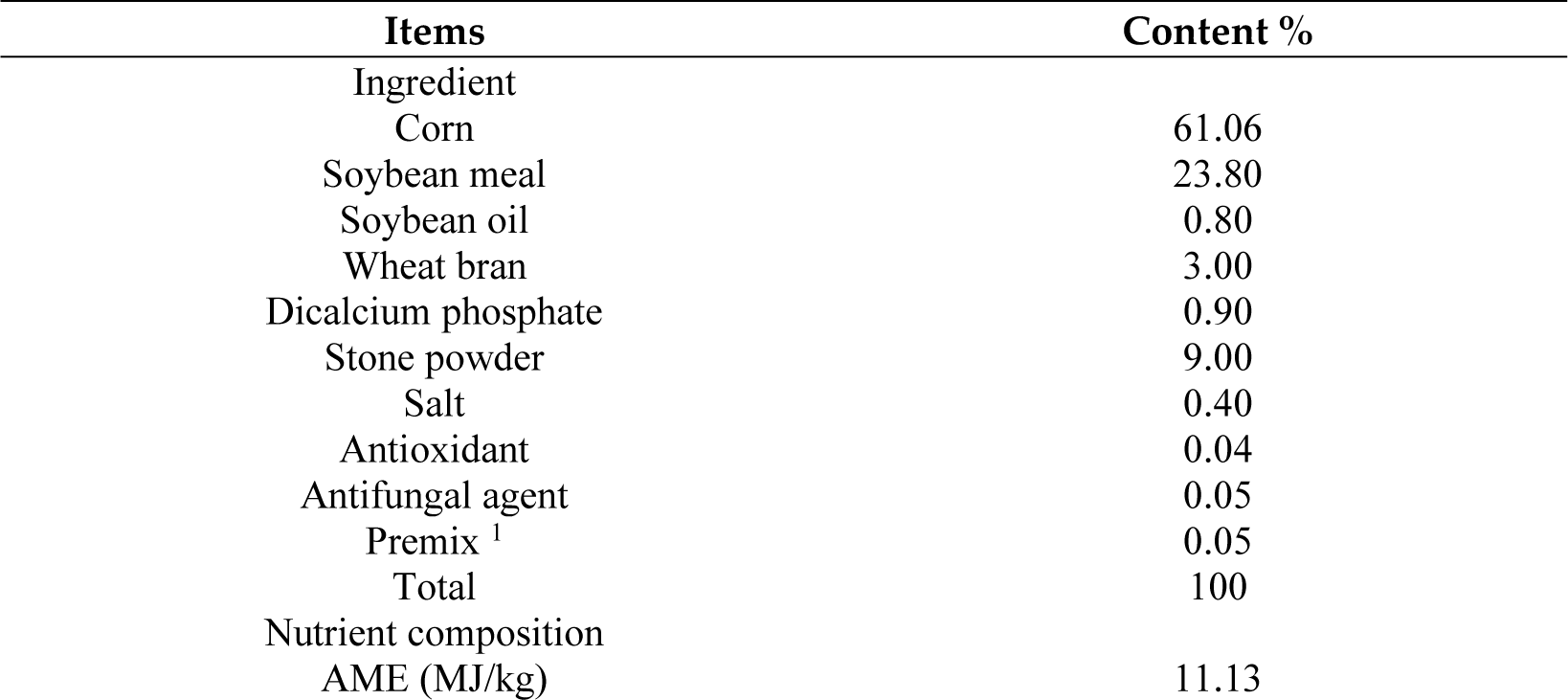
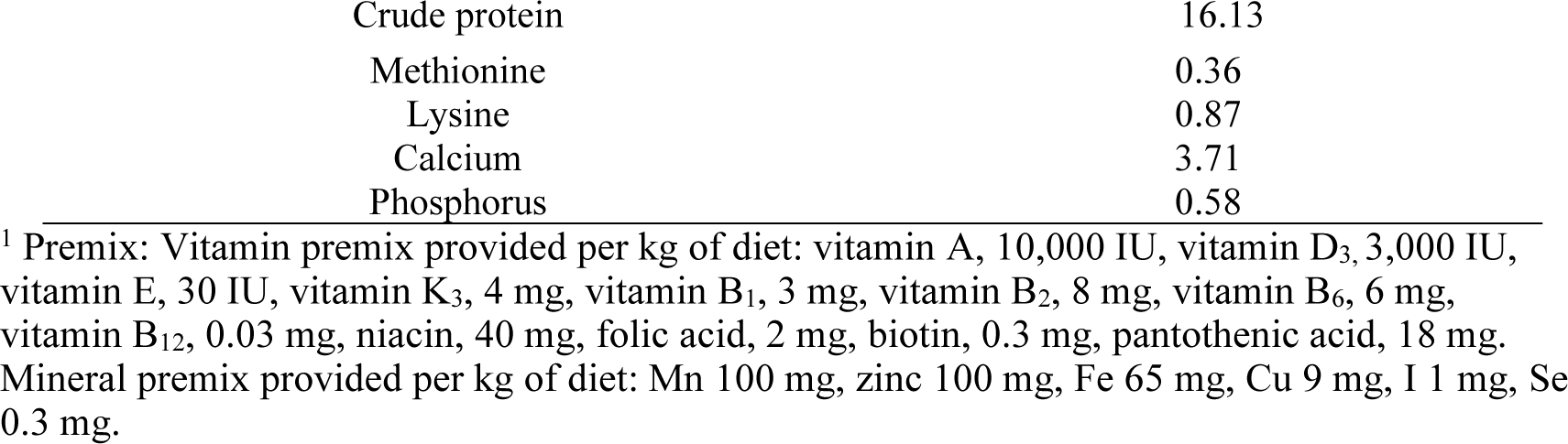
Composition and nutrient levels of the basal diet (as-fed basis, %)

| Items | Content % |
| --- | --- |
| Ingredient |  |
| Corn | 61.06 |
| Soybean meal | 23.80 |
| Soybean oil | 0.80 |
| Wheat bran | 3.00 |
| Dicalcium phosphate | 0.90 |
| Stone powder | 9.00 |
| Salt | 0.40 |
| Antioxidant | 0.04 |
| Antifungal agent | 0.05 |
| Premix <sup>1</sup> | 0.05 |
| Total | 100 |
| Nutrient composition |  |
| AME (MJ/kg) | 11.13 |
| Crude protein | 16.13 |
| Methionine | 0.36 |
| Lysine | 0.87 |
| Calcium | 3.71 |
| Phosphorus | 0.58 |

<sup>1</sup> Premix: Vitamin premix provided per kg of diet: vitamin A, 10,000 IU, vitamin D<sub>3</sub>, 3,000 IU, vitamin E, 30 IU, vitamin K<sub>3</sub>, 4 mg, vitamin B<sub>1</sub>, 3 mg, vitamin B<sub>2</sub>, 8 mg, vitamin B<sub>6</sub>, 6 mg, vitamin B<sub>12</sub>, 0.03 mg, niacin, 40 mg, folic acid, 2 mg, biotin, 0.3 mg, pantothenic acid, 18 mg. Mineral premix provided per kg of diet: Mn 100 mg, zinc 100 mg, Fe 65 mg, Cu 9 mg, I 1 mg, Se 0.3 mg.

At the end of the experimental period (70 weeks), outliers that were more than 3 standard deviations (SD) away from the mean in terms of egg weight (EW), body weight (BW), and total feed intake (TFI) were removed. As a result, a total of 1192 hens from the base population were identified for the subsequent analysis of the correlation between RFI and production performance as well as feed efficiency.

From the base population of 1192 hens, the lowest 10 hens and the highest 10 hens were chosen according to their remaining RFI value (Defined as the LRFI group and the HRFI group), while ensuring no differences in preslaughter body weight and egg production performance. Total 20 hens (10 hens/each group) were used for analysis of egg quality, serum, liver biochemistry, body measurements, and intestinal morphological structure. Due to sample loss, 8 hens were finally retained from each group of 10 hens for transcriptome, metabolome, and microbiome analysis. The whole test process was carried out in strict accordance with the scheme approved by the Animal Welfare Committee of China Agricultural University (AW62801202-1-1).

### 2.2 Laying performance and egg quality determination

The egg number (EN), EW, and mortality were recorded daily. The egg laid (44, 57, and 70 weeks of age), BW at 43 weeks by electronic scales (accuracy to 0.01 g). First egg period (FEP), first egg weight (FEW) and first body weight (FBW) were recorded.

At 70 weeks of age, egg quality was recorded. The egg shape index (ESI) was calculated by the formula: egg length/egg width; eggshell color (ESC) was measured by the fast and easy-to-use CIE L*a*b* system^[21]^; EW was obtained with an electronic balance (YP601N, Qinghai Co., Ltd., Shanghai, China)^[22]^.

### 2.3 Calculation of feed efficiency

Feed consumption, BW, and egg mass (EM) were divided into two measurement cycles from 30-31 and 67-69 weeks of age. Separate metal feed troughs were used to provide mash feed to each hen. Feed is added by hand every day after the weighing tank. The weight of each hen was measured using an electronic scale at the beginning and end of the feeding trial. Feed intake (FI) was calculated weekly, EW and EN were recorded daily. The TFI of individuals was calculated based on the feed intake data. Body weight gain (BWG) is the difference in weight between the start and end of a feeding test. The FI for each interval and daily feed intake (DFI) for each hen were calculated. Daily egg mass (DEM) is calculated as the product of the average egg weight (AEW) on the test day of the total number of eggs. The metabolic body weight (MBW: mean body weight from 30-31 or 69-70 weeks of age), FCR and RFI were calculated. The FCR is calculated as the ratio of DFI and DEM. The RFI is estimated according to the following formula^[23]^.

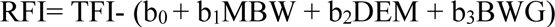

where the TFI encompassed that from 30-31 and 69-71 weeks of age, b_0_ is the intercept, and b_1_, b_2_ and b_3_ are partial regression coefficients.

### 2.4 Characteristics of digestive organs and body measurement

At the end of the experimental period (70 weeks), samples were obtained from 20 chickens (10/group). The following digestive organs in the abdominal cavity were measured with an electronic balance with an accuracy of 0.01g, including liver weight, pancreatic weight, proventriculus weight, gizzard weight (contents and surrounding fat removed), duodenum weight, jejunum weight, ileum weight and cecum weight. The length of each intestinal segment was measured with a measuring tape to the nearest 1 cm, including duodenum length, jejunum length, ileum length and cecum length (total length on both sides).

The body measurement was measured by tape. The contents of the measurement mainly included live weight before slaughter (LWBS), body slope length (BSL), keel length (KL), chest width (CW), chest depth (CD), shank length (SL), and shank girth (SG), which were measured by *performance terminology and measurement for poultry* (NY/T 823-2020).

### 2.5 Determination of serum biochemical indexes

Before slaughtering (70 weeks of age), blood samples were obtained from 20 chickens (10/group). All blood samples were maintained on ice until centrifugation at 3,000 * *g* for 15 min at 4 °C^[24]^. After the blood cells had completely settled to the bottom of the tube, the upper serum was transferred into a centrifuge tube with a pipette at −80°C for further analysis. Serum samples were collected to determine the concentrations of hormones, including insulin (INS), leptin (LEP), adrenocorticotropic hormone (ACTH), and growth hormone (GH), measured by ELISA kits (Kehua Bioengineering Co., Ltd, Shanghai, China). Another serum sample was used to analyze lipid profiles, including total cholesterol (TC), triglycerides (TG), low-density lipoprotein cholesterol (LDL-C), high-density lipoprotein cholesterol (HDL-C), alanine aminotransferase (ALT), and aspartate aminotransferase (AST), measured by an automatic biochemical instrument (Kehua, KHB-1280, Shanghai, China).

### 2.6 Hematoxylin-eosin staining and Oil Red O staining

The liver was fixed with 4% paraformaldehyde, embedded in paraffin wax and cut into 4-5 μm sections, and the fresh liver tissue was fixed with fixation liquid for more than 24 hours. H&E and Oil Red O staining were performed according to standard protocols to assess the occurrence of lesions and fatty deposits in the liver^[24,25]^.

### 2.7 RNA extraction, transcription data analysis and quantitative real-time PCR

Total RNA was extracted from the liver tissue using TRIzol Reagent according to the manufacturer’s instructions. Only high-quality RNA samples (OD260/280=1.8∼2.2) were used to construct the sequencing library. According to Illumina’s protocol, cDNA libraries were constructed using RNA from high quality liver tissues using the TruSeq™ Strand Total RNA Library Preparation Kit reagent (Mayo Biomedical Technology Co., Ltd., Shanghai, China)^[26]^. All processes were performed in accordance with the manufacturer’s instructions and then sequenced on the Illumina HiSeq platform at Majorbio Biopharm Technology Co., Ltd. (Shanghai, China).

To identify differentially expressed genes (DEGs) between two different RFI groups, the raw counts were statistically analyzed using DESeq_2_ software. The DEGs with|log_2_(fold change)|≥1 and *p-*values < 0.05 were considered DEGs. The GOG database (http://eggnog.embl.de/) and the KEGG database (http://www.genome.jp/kegg/) were used for DEG annotations^[27]^.

After the total RNA was extracted, reverse transcription of mRNA and *real-time quantitative PCR* (qRT–PCR) were conducted by using a BeyoRT™II cDNA synthesis kit and BeyoFast™ SYBR Green qPCR Mix (Beyotime Biotechnology Co., Ltd., ShangHai, China) were conducted in duplicate in a CFX-96 real-time PCR detection system (Bio-Rad Laboratories, Hercules, CA, USA). Using β-actin as the internal reference gene, the primer sequences are shown in supplemental information, synthesized by Shanghai Bioengineering Company. Each sample was repeated 3 times, and the 2^−ΔΔCt^ method was used to calculate the relative expression of the mRNA of the target gene^[28]^. The protocol for all genes was as follows: 95 °C for 15 min; 40 cycles of 95 °C for 10 s and 60 °C for 30 s.

### 2.8 16S rRNA sequencing in cecum microbiota

Total microbial genomic DNA was extracted from cecum content samples using the E.Z.N.A.® soil DNA Kit (Omega Biotek, Norcross, GA, U.S.) according to the manufacturer’s instructions. Checking the concentration and purity of the DNA sample using a NanoDrop 2000 spectrophotometer. The highly variable region V3-V4 of the bacterial 16S rRNA gene was amplified with primers for 338F (5’- ACTCCTACGGGAGGCAGCAGCAGCAG-3’) and 806R (5’-GGACTACHVGGGTWTCTAAT-3’). At Majorbio Company, paired-end sequencing of amplicons was performed using the Illumina MiSeq sequencing platform and PE300 chemistry.

The data were normalized and transformed, and then correlation analysis, cluster analysis, and statistical analysis were used to identify differentially abundant metabolites (DEMs), and heatmaps were drawn.

### 2.9 Untargeted metabolomics analysis

The instrument platform for liquid chromatography–mass spectrometry (LC-MS) analysis was the UHPLC-Q exactive system from Thermo Fisher Scientific by Majorbio Biopharm Technology Co., Ltd (Shanghai, China). After mass detection was completed, the raw LC-MS data were preprocessed by Progenesis Qi (Waters Corporation, Milford, USA) software and the matrix was exported in CSV format. The liver metabolites were searched and identified, and the main databases were HMDB (http://www.hmdb.ca/), Metlin (https://metlin.scripps.edu/) and the Majorbio database. These liver metabolites were annotated using the KEGG database (http://www.genome.jp/kegg/). The R package (Version1.6.2) software was used to screen the DEMs, and metabolites with a *p-*value < 0.05 and VIP >1 were considered DEMs^[29, 30]^.

## 3. Results

### 3.1 Descriptive Statistics of Chicken Phenotypes

As shown in Table 2, the descriptive statistics of all phenotypic traits, including the mean, SD, coefficient of variation (CV), minimum, and maximum, are summarized. From all experimental chicken populations (N=1192), the mean values of EM_30-31_, MBW_30-31_, FI_30-31_, FCR_30-31_, and RFI_30-31_ were 766.24 g, 296.32 g, 1700.94 g/d, 2.24 g/g, and 1.94 g/d, respectively. Notably, the CV of FI was greater than 10%, indicating a large phenotypic variation in these traits, and the RFI was approximately equal to zero because it represented the residuals of the linear model.

**Table 2.** Descriptive statistics for phenotypes of pedigreed hens.

| Traits<sup>1</sup> | N | Mean | SD | CV (%) | Maximum | Minimum |
| --- | --- | --- | --- | --- | --- | --- |
| EN<sub>44</sub> | 1192 | 155.57 | 10.00 | 6.45 | 184 | 118 |
| EN<sub>57</sub> | 1192 | 242.54 | 16.61 | 6.85 | 277 | 132 |
| EM<sub>30-31</sub>(g) | 1192 | 766.24 | 96.71 | 12.62 | 1025.56 | 471.96 |
| BW<sub>30</sub>(g) | 1192 | 1979.34 | 166.45 | 8.41 | 2536.04 | 1525.01 |
| BW<sub>31</sub>(g) | 1192 | 1955.34 | 149.39 | 7.64 | 2475.38 | 1500.04 |
| MBW<sub>30-31</sub>(g) | 1192 | 296.32 | 18.70 | 6.31 | 357.36 | 244.03 |
| FI<sub>30-31</sub>(g/d) | 1192 | 1700.94 | 146.98 | 8.64 | 2135.00 | 1237.28 |
| FCR<sub>30-31</sub>(g/g) | 1192 | 2.24 | 0.31 | 13.84 | 3.54 | 1.31 |
| RFI<sub>30-31</sub>(g/d) | 1192 | 1.94 | 139.33 | - | 425.99 | -518.53 |

<sup>1</sup> N, Number of hens with phenotypic value; EN<sub>44/57</sub>, total egg number at 44/57 weeks of age; EM<sub>30-31</sub>, total egg mass from 30 to 31 weeks of age; BW<sub>30/31</sub>, body weight at 30/31 weeks of age; MBW<sub>30-31</sub>, mean body weight from 30 to 31 weeks of age; FI<sub>30-31</sub>, feed intake from 30 to 31 weeks of age; FCR<sub>30-31</sub>, feed conversion ratio from 30 to 31 weeks of age; RFI<sub>30-31</sub>, residual feed intake from 30 to 31 weeks of age.

The correlation coefficient between growth performance and feed efficiency traits is shown in Fig. 1. By calculating the correlation between RFI_30-31_ and BW_43_, EN_27_, EN_44_, EN_57_, FI_30-31_, EW_30-31_, FEP, FEW, FBW in the experimental chickens, it was found that there was a significant strong correlation between RFI and FI (R=0.94 >0.3), while the correlation with other production performance indicators was weak (R<0.1, negligible). Referring to the literature related to the correlation with RFI, the correlation results of this experimental chicken population are reasonably consistent^[31]^.

**Fig. 1.**
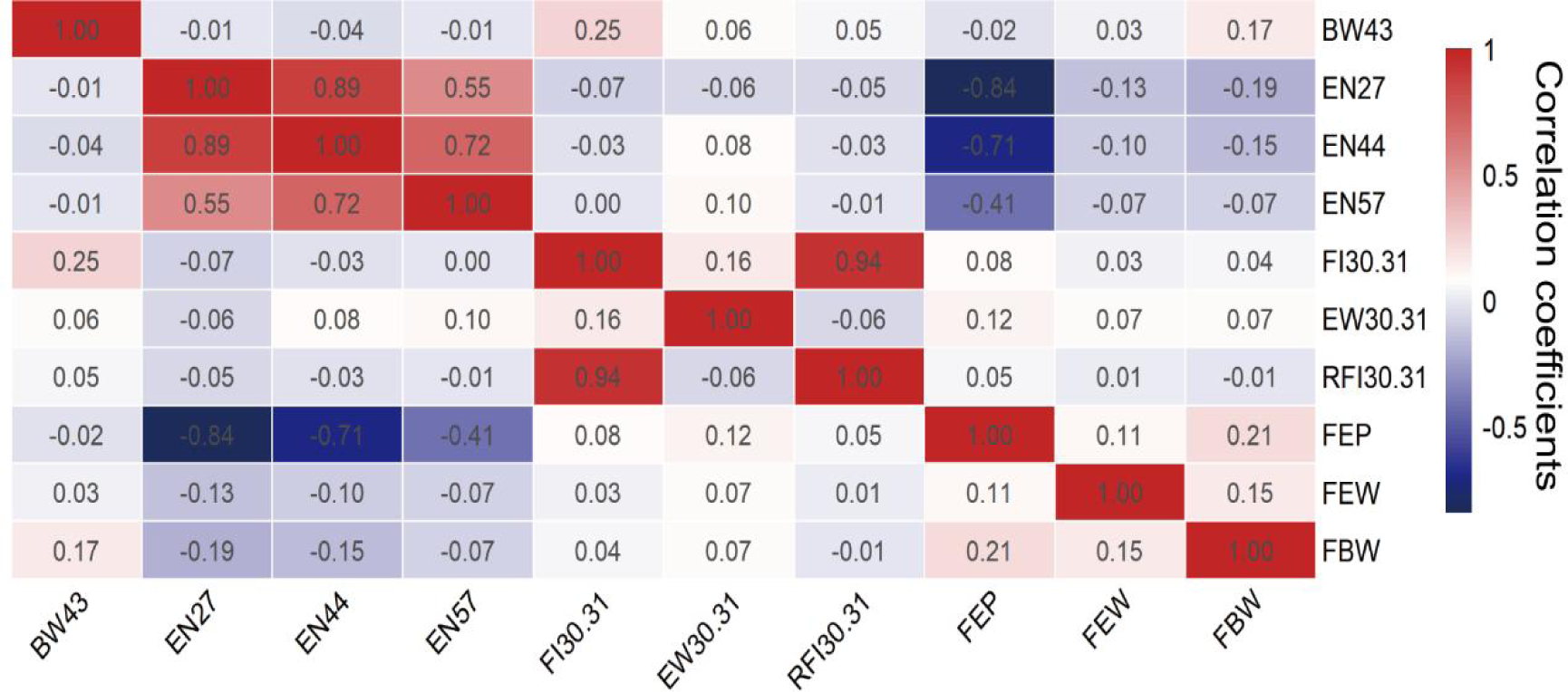
The correction of RFI and production performance of pedigreed hens. BW_43,_ body weight at 43 weeks of age; EN_27/44/57_, total egg number at 27/44/57 weeks of age; FI_30-31,_ feed intake from 30 to 31 weeks of age; EW_30-31_, egg weight from 30 to 31 weeks of age; RFI_30-31,_ residual feed intake from 30 to 31 weeks of age. FEP, first egg period; FEW, first egg weight; FBW, first body weight of hens; The red and blue gradients indicate positive and negative correlation coefficients, respectively. The redder or bluer the block color is, the greater the correlation coefficient. The whiter the area is, the smaller the correlation coefficient.

At 70 weeks of age, chickens (10/group) were selected from the base experimental population for subsequent analysis. As shown in Table 3, the FI, FCR and RFI of the HRFI group were higher than those of the LRFI group (*P* < 0.05). The mean RFI of the high and low RFI groups was 93.19 g/d and −117.94 g/d, respectively. Moreover, there was no significant difference in EN_70,_ FEP, FBW, AEW_67-69,_ MBW_67-69_ between the high and low RFI groups (*P>0.05*).

**Table 3.** Production performance of laying hens.

| Traits<sup>1</sup> | LRFI | HRFI |
| --- | --- | --- |
| EN<sub>70</sub> | 313.20±10.01 | 313.90±4.61 |
| FEP(d) | 139.00±5.66 | 141.20±5.39 |
| FBW(g) | 1768.90±62.82 | 1766.20±61.11 |
| AEW<sub>67-69</sub>(g) | 57.61±2.78 | 59.16±2.80 |
| MBW<sub>67-69</sub>(g) | 294.79±18.57 | 292.96±15.71 |
| FI<sub>67-69</sub>(g) | 2305.28±51.50<sup>b</sup> | 2513.94±39.93<sup>a</sup> |
| FCR<sub>67-69</sub>(g:g) | 2.37±0.35<sup>b</sup> | 2.82±0.41<sup>a</sup> |
| RFI<sub>67-69</sub>(g/d) | -117.94±25.72<sup>b</sup> | 93.19±20.22<sup>a</sup> |

<sup>1</sup> EN<sub>70</sub>, total egg number at 70 weeks of age; FEP, first egg period; FBW, first body weight of hens; AEW<sub>67-69</sub>, average egg weight from 67 to 69 weeks of age; MBW<sub>67-69</sub>, mean body weight from 67 to 69 weeks of age; FI<sub>67-69</sub>, feed intake from 67 to 69 weeks of age; FCR<sub>67-69</sub>, feed conversion ratio from 67 to 69 weeks of age; RFI<sub>67-69</sub>, residual feed intake from 67 to 69 weeks of age. Values are presented as the mean and SD ( $n = 10$ , 67-69 weeks of age), and different letters indicate statistically significant differences ( $P < 0.05$ ).

Egg quality serves as an important economic indicator for evaluating the superiority or inferiority of eggs. As shown in Table 4, at 70 weeks of age, the egg quality was tested. As expected, there were no significant differences observed in terms of ESI, ESC and EW between the HRFI and LRFI groups *(P > 0.05*). These results also indicated that different RFI groups of laying hens only exhibited a significant difference in feed efficiency, while there were no significant differences in terms of production performance and egg quality.

**Table 4.** Egg quality of laying hens.

| Traits<sup>1</sup> | LRFI | HRFI |
| --- | --- | --- |
| ESI | 1.32±0.05 | 1.32±0.04 |
| ESC-L | 62.33±6.28 | 62.69±1.72 |
| ESC-A | 14.75±3.89 | 15.34±0.98 |
| ESC-B | 25.80±3.55 | 27.57±1.57 |
| EW(g) | 58.11±3.62 | 58.53±2.75 |

<sup>1</sup>ESI, egg shape index; ESC-L, eggshell color-L; ESC-A, eggshell color-A; ESC-B, eggshell color-B; EW, egg weight. Values are presented as the mean and SD ( $n = 10$ , 70 weeks of age).

Body size is the simplest trait that can be directly measured, which simplifies the selection and mating process for indirectly related traits with strong correlations. The differences between the HRFI and LRFI groups at 70 weeks in terms of body measurements are provided in Table 5. Measurements revealed that the CW of the LRFI group of hens was significantly higher than that of the HRFI group (*P < 0.05*), while there were no significant differences observed between the two groups in terms of LWBS, BSL, KL, CD, SL, and SG (*P > 0.05*).

**Table 5.** Body size traits of laying hens.

| Traits<sup>1</sup> | LRFI | HRFI |
| --- | --- | --- |
| LWBS(g) | 2088.50±173.88 | 2059.50±138.63 |
| BSL(cm) | 19.37±0.53 | 19.20±0.75 |
| KL(cm) | 12.30±0.75 | 12.27±0.77 |
| CW(cm) | 81.75±2.67<sup>a</sup> | 78.34±4.17<sup>b</sup> |
| CD(cm) | 115.36±4.15 | 114.30±5.42 |
| SL(cm) | 8.81±0.29 | 8.65±0.27 |
| SG(cm) | 4.92±0.27 | 4.75±0.43 |

<sup>1</sup>LWBS, liver weight before slaughter; BSL, body slope length; KL, keel length; CW, chest width; CD, chest depth; SL, shank length; SG, shank girth. Values are presented as the mean and SEM ( $n = 10$ , 70 weeks of age), and different letters indicate statistically significant differences ( $P < 0.05$ ).

### 3.2 Characteristics of Digestive Organs

The length and weight of the digestive organs in laying hens are meaningful in evaluating their digestive function and feed utilization efficiency. After slaughter, the length and weight of the digestive organs in the two groups are shown in Figure 2A-B, 2E. From the results, it can be observed that in terms of organ weight, the gizzard, ileum and cecum weight of hens in the LRFI group were significantly higher than those of in the HRFI group *(P < 0.05)*. Regarding organ length, the duodenum and cecum lengths of the LRFI group were significantly higher than those of the HRFI group *(P < 0.05)*. Besides, there were no significant differences observed between the two groups in terms of liver weight, pancreatic weight, ventriculus weight, duodenum weight, jejunum weight, jejunum length, or ileum length (*P>0.05*).

**Fig. 2.**
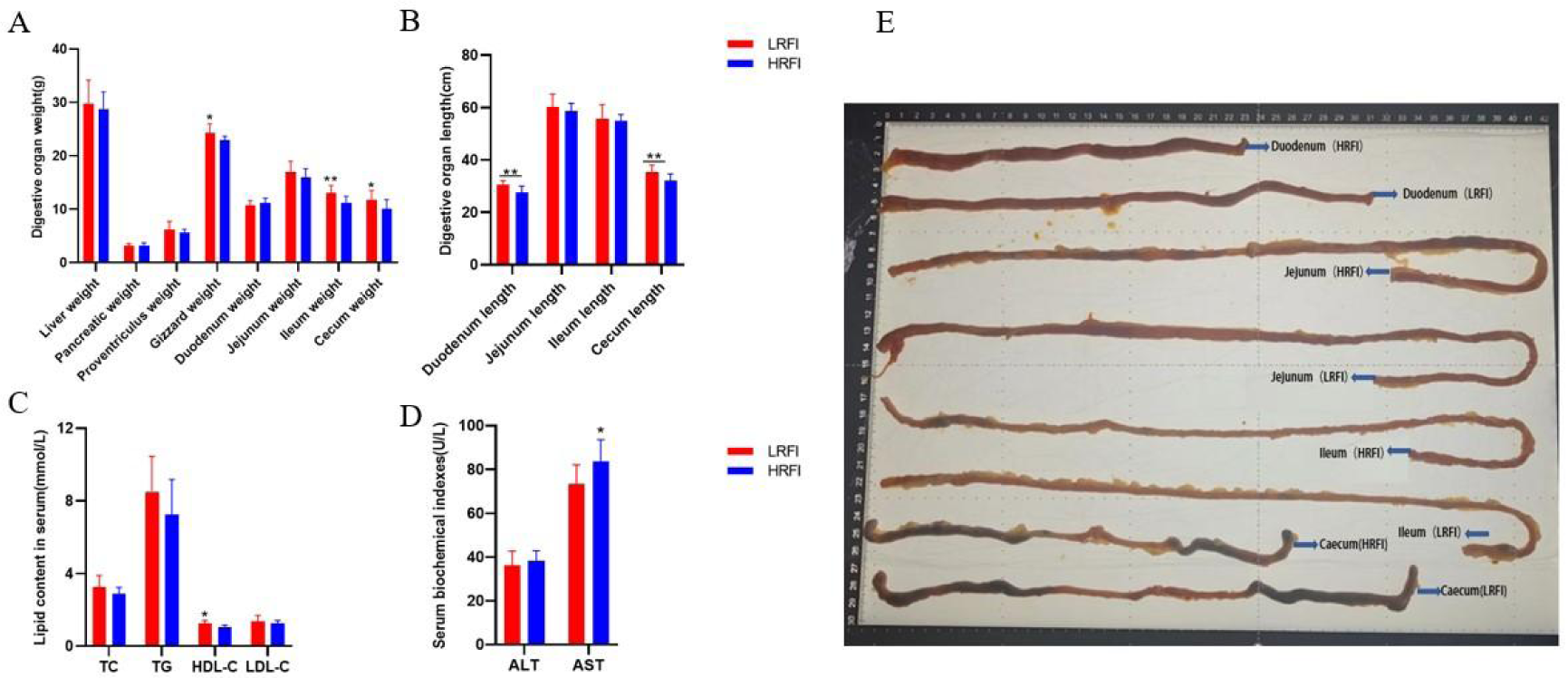
Different RFI groups on (A) digestive organ weight; (B) digestive organ length; (C) TC, TG, HDL-C, LDL-C content in serum; (D) ALT, AST content in serum. (E) Representative images of poultry intestines at sacrifice. Data are represented as the means ± SD (n = 10). Asterisks (* or **) indicate a significant difference at *p < 0.05* or *p < 0.01*, respectively.

### 3.3 Plasma hormones of laying hens

Serum hormones and physicochemical indicators can effectively reflect the changes in feed intake, nutritional requirements, physiological status and metabolic levels of laying hens. The differences in hormone levels between the HRFI and LRFI hens are illustrated (Fig. 2 C-D and Table 6). Compared with the HLRI group, the serum ACTH and AST contents in the LRFI group were significantly decreased and the serum HDL-C contents were significantly increased (*P < 0.05*).

**Table 6.**
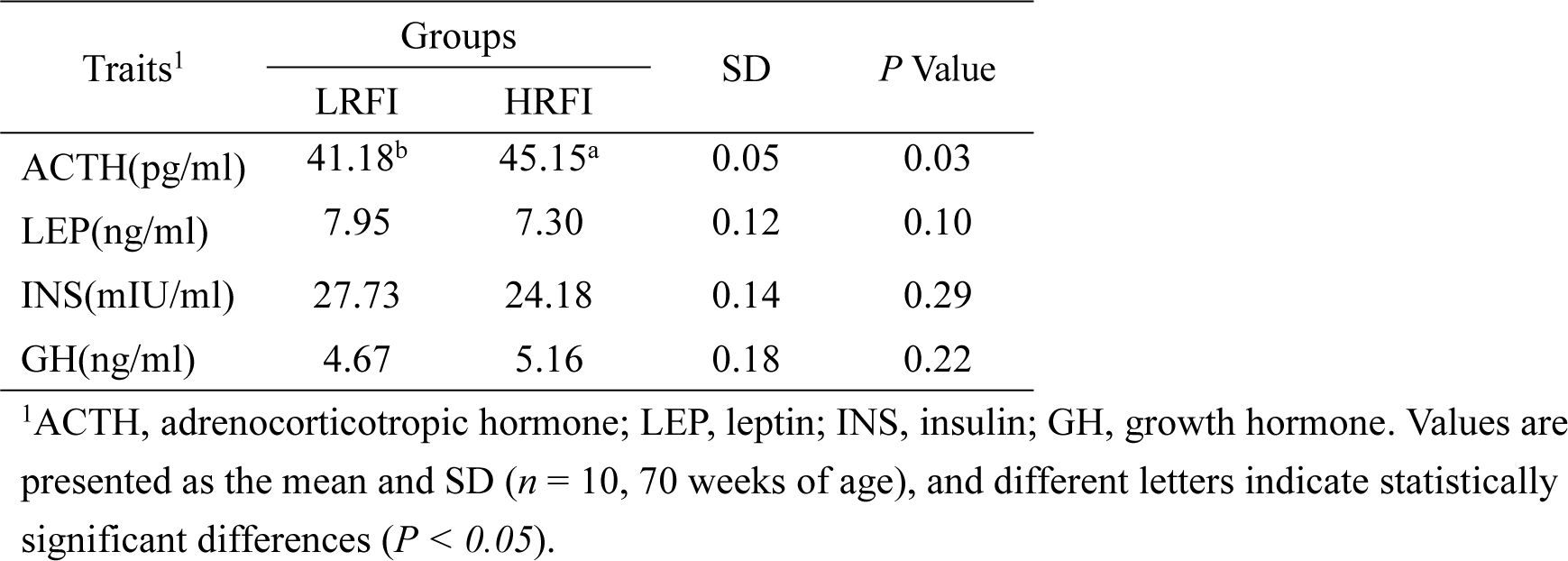
Hormone levels in the serum of laying hens.

| Traits<sup>1</sup> | Groups |  | SD | P Value |
| --- | --- | --- | --- | --- |
|  | LRFI | HRFI |  |  |
| ACTH(pg/ml) | 41.18<sup>b</sup> | 45.15<sup>a</sup> | 0.05 | 0.03 |
| LEP(ng/ml) | 7.95 | 7.30 | 0.12 | 0.10 |
| INS(mIU/ml) | 27.73 | 24.18 | 0.14 | 0.29 |
| GH(ng/ml) | 4.67 | 5.16 | 0.18 | 0.22 |

<sup>1</sup>ACTH, adrenocorticotrophic hormone; LEP, leptin; INS, insulin; GH, growth hormone. Values are presented as the mean and SD ( $n = 10$ , 70 weeks of age), and different letters indicate statistically significant differences ( $P < 0.05$ ).

### 3.4 Histopathological evaluation of hen liver

Figure 3 presents the results observed under electron microscopy after liver tissue staining. H&E staining results show that the liver cells in the LRFI group are tightly arranged with normal morphological structures. The cell nuclei are clearly visible, and there are no significant pathological changes. On the other hand, liver cells in the HRFI group exhibited slight atrophy, increased intercellular spaces, indicating a tendency toward vacuolation. Additionally, compared to the HRFI group, Oil Red O staining indicated that the liver cells in the HRFI group showed a certain degree of lipid accumulation, with significantly increased lipid droplets and slightly blurred morphological structures.

**Fig. 3.**
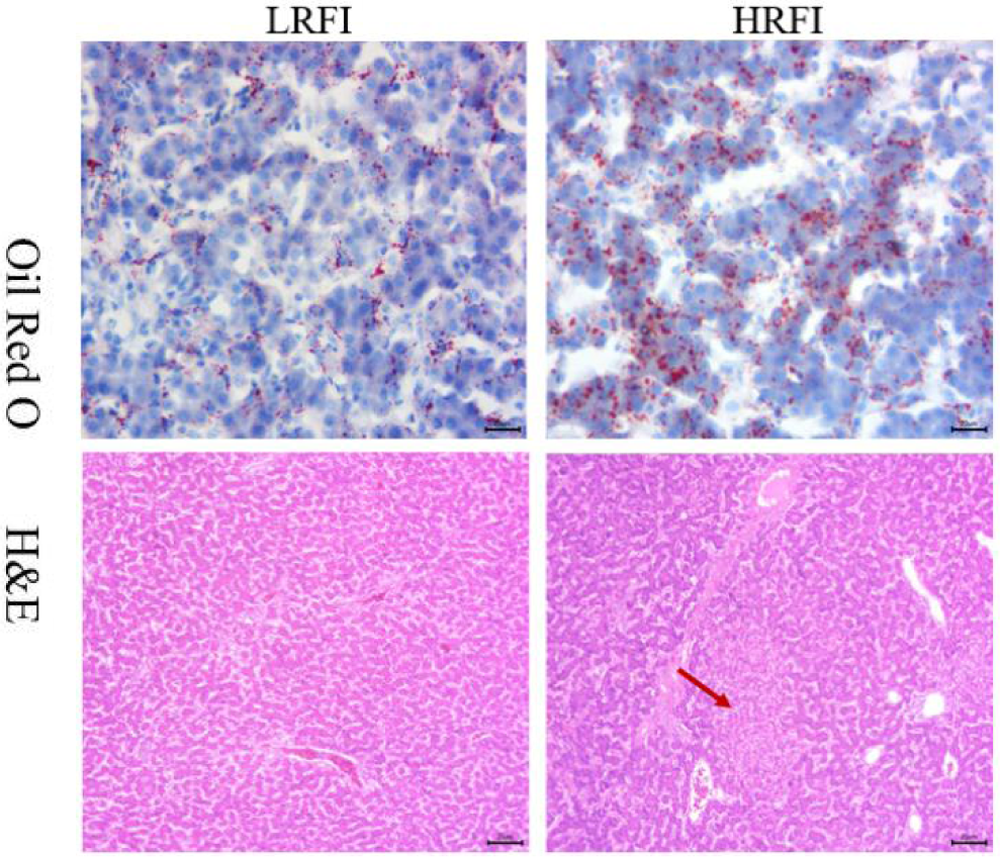
Liver tissue structure of different RFI groups. H&E staining, 200× magnification; Oil Red O staining, 400× magnification (Red arrows indicate cytoplasmic vacuolation.).

### 3.5 Identification of target genes in the liver

To reveal the transcriptional changes in the LRFI and HRFI groups, RNA-Seq analysis was performed. After sequencing on the Illumina platform (PE libraries with a read length of 2×150 bp), the raw sequences obtained from 16 liver samples ranged from 41,738,168 to 43,661,442. Following quality control, the range of clean reads was 41,260,626 to 42,784,452. The Q_20_ and Q_30_ values for all samples exceeded 96% and 91%, respectively. The GC content ranged from 48.42% to 51.55%, and the error rate was less than 0.03%. These results indicate that the filtered sequences obtained from this sequencing experiment are suitable for subsequent bioinformatics analysis (Supplementary Table). HISAT2 software (http://ccb.jhu.edu/software/hisat2/index.shtml) was employed to align the filtered sequences after quality control to the chicken (*Gallus gallus*) reference genome (GRCg6a, http://asia.ensembl.org/Gallus_gallus/Info/Index). More than 93% of the sequences were successfully aligned to the reference genome, with 90.44% to 91.88% of the sequences uniquely mapped to the genome. This indicates the high accuracy and reliability of the sequencing data obtained in this experiment. Subsequent analyses were based on the uniquely mapped reads (Supplementary Table).

A Venn diagram showed that 9900 genes were enriched in the two groups, and each group contained 237 and 872 DEGs, respectively (Fig. 4A). The results (Fig. 4B) showed that 101 DEGs out of the 17,870 DEGs were downregulated (DDEGs) in the LRFI vs HRFI group, while 107 DEGs were upregulated (UDEGs). Transcriptional changes in the LRFI and HRFI groups were investigated using RNA-Seq analysis. After aligning the filtered sequences to the chicken reference genome, a total of 268 DEGs were identified between the two groups based on a threshold of *p<0.05* and fold change >2.0 by using t-tests. The DEGs from all comparison groups were combined to form a set of DEGs. Cluster analysis was performed on this gene set to group genes with similar expression patterns and identify genes with the same expression trends (Fig. 4C).

**Fig. 4.**
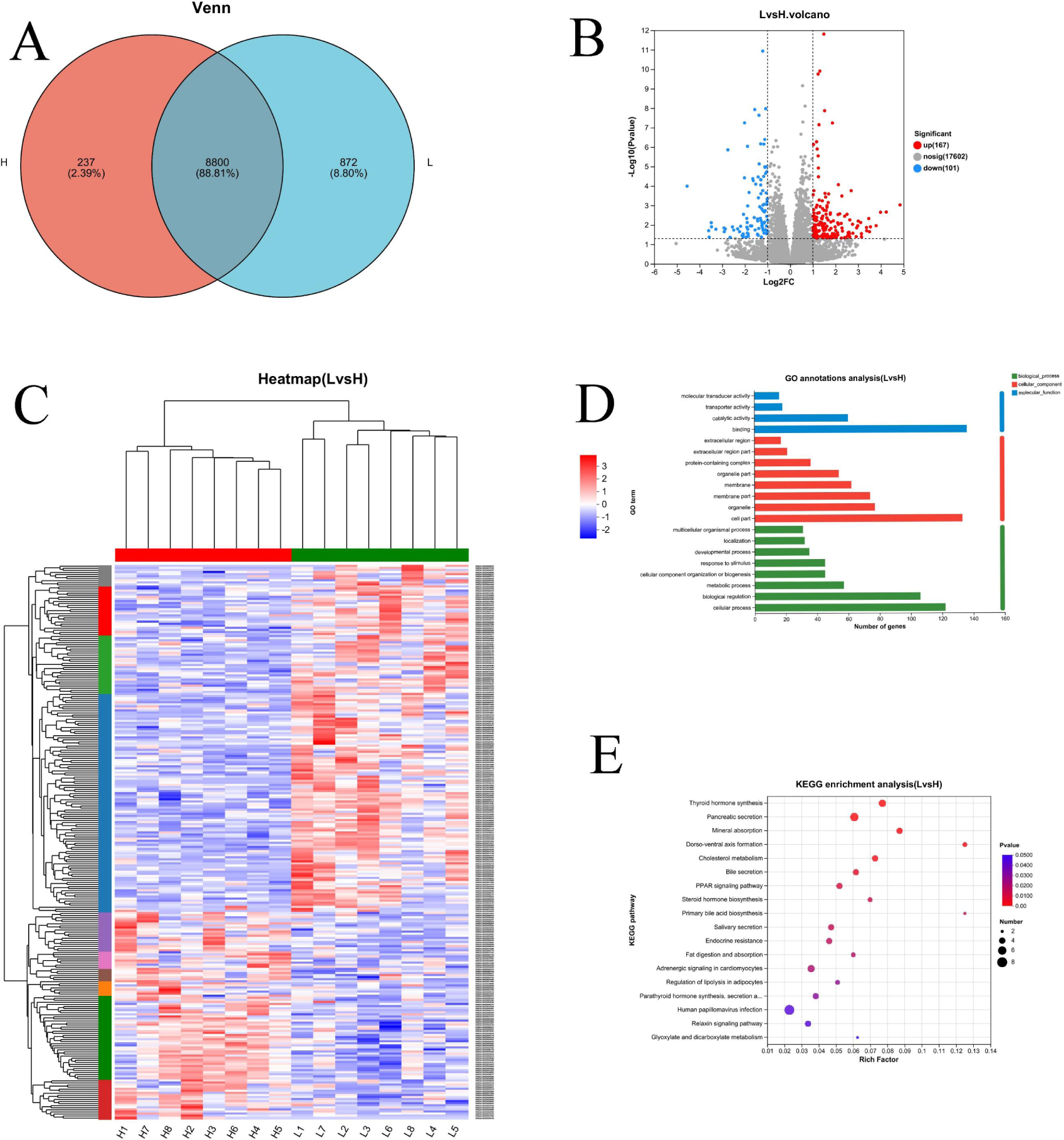
RNA-seq data analysis of liver in the LRFI and HRFI groups (n = 8 per group). (A) Venn diagram of DEGs; (B) Volcano plot; (C) DEGs expression in cluster heatmap; (D) GO annotation analysis; (E) KEGG pathway enrichment analysis.

The DEG functions were classified using GO enrichment analysis. The findings revealed that the top 20 enriched GO terms that were derived after comparing the abovementioned 3 groups belonged to the 3 GO categories of CC, BP, and MF (Fig. 4D). GO functional annotation analysis was performed on the DEGs identified in this experiment. The major annotated cellular components include: cell part, organelle, organelle part, membrane, membrane part, protein-containing complex; DEGs involved in molecular functions significantly enriched in binding, catalytic activity, and transporter activity; DEGs involved in biological processes significantly enriched in cellular process, biological regulation, metabolic process, cellular component organization or biogenesis, response to stimulus. The maximum number of DEGs was enriched in the ‘cellular process’ (122 DEGs), followed by ‘cell part’ (133 DEGs) and ‘binding’ (136 DEGs).

The DEGs were then used for KEGG analysis (Fig. 4E). A total of 18 KEGG pathways were significantly enriched, with the five most significant pathways including ‘Thyroid hormone synthesis’, ‘Pancreatic secretion’, ‘Mineral absorption’, Dorsoventral axis formation and Cholesterol metabolism. Finally, 30 up and downregulated DEGs were significantly enriched in these significant KEGG pathways (Supplemental information). The DEGs in the beforementioned KEGG pathways may play a role in regulating energy, hormones, substance and nutrient absorption in hens during the late laying period. Therefore, they can be considered candidate genes involved in regulating feeding behavior.

To validate the RNA-Seq data, 6 DEGs related to RFI were selected and analyzed by qRT‒PCR. The results of analysis showed similar expression trends as RNA-Seq analysis, which indicated the reproducibility and reliability of the RNA-Seq data (Fig. 5).

**Fig. 5.**
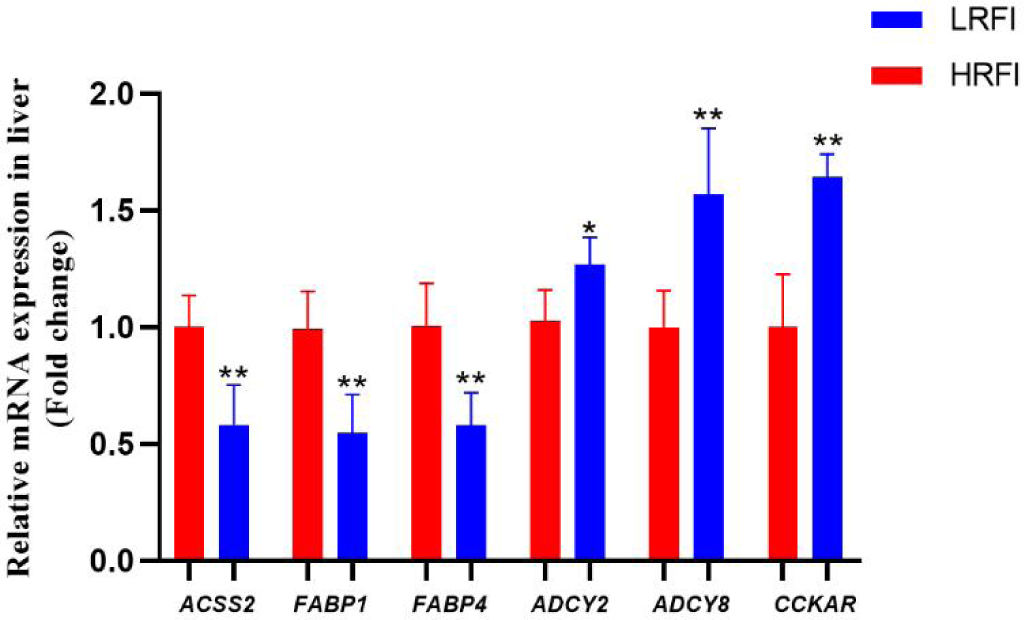
qRT‒PCR validation in the livers of the LRFI and HRFI groups. Data are represented as the mean ± SD (n = 8). Asterisks (* or **) indicate a significant difference at *p < 0.05* or *p < 0.01*, respectively.

### 3.6 Cecal microbial composition of laying hens

The similarity and overlap of OTUs between the LRFI and HRFI groups are summarized in the Venn diagram (Fig. 6A). The results implied that the two groups shared 1045 OTUs, while some OTUs were unique to specific groups (282 for the LRFI group and 233 for the HRFI group), showing that the intestinal bacteria exerted a certain effect on the OTU composition of the two groups. The cecum microbial alpha diversity, including the Ace, Chao, Shannon and Sobs indices of the HRFI group was lower (*P < 0.05*) than that of the LRFI group. In addition, PCA exhibited a significant difference in species composition between the two groups, showing the importance of investigating RFI from the perspective of microorganisms (Fig. 6B). These results showed that compared with the HRFI group, the bacterial diversity and richness of the LRFI group changed significantly.

**Fig. 6.**
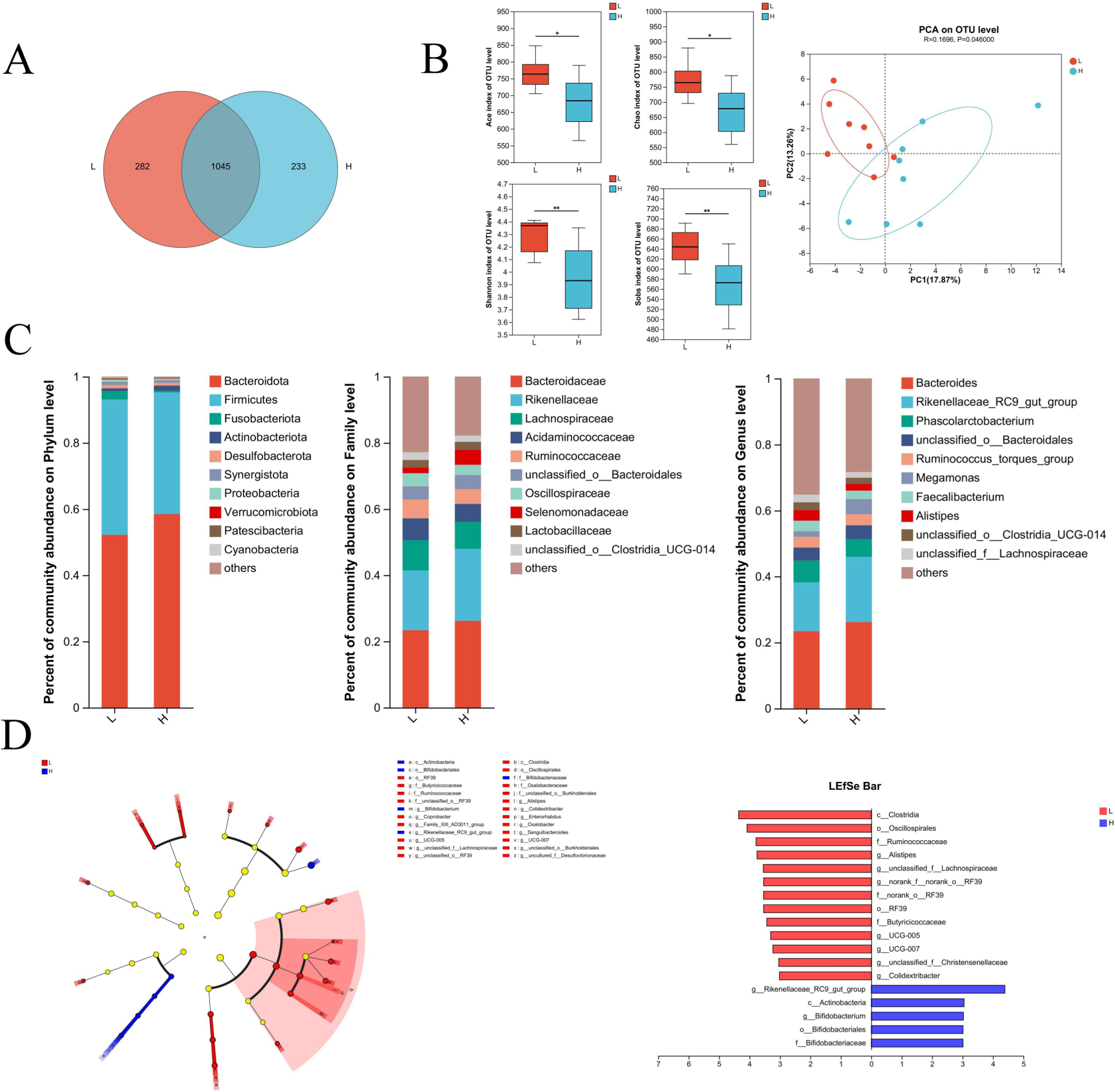
Cecal microbial composition in the LRFI and HRFI groups (n = 8 per group). (A) Venn diagram; (B) Alpha and Beta diversity (PCA) by T test; (C) Alterations in microbiota at the phylum, family, and genus levels; (D) Levels and biomarkers found by linear discriminant analysis effect size (LEfSe) and LDA scores. Species with significant differences that have an LDA score >3 are presented. Data are presented as the means ± SD (*\*P < 0.05; **P < 0.01*).

As illustrated in Fig. 6C, the dominant phyla in the LRFI and HRFI groups were *Bacteroidota* and *Firmicutes.* At the phylum level, the LRFI group showed an increase in relative abundance of *Firmicutes* and a decrease in relative abundance of *Bacteroidetes*. At the family and genus levels, *Bacteroidaceae/Bacteroides* and *Rikenellaceae/Rikenellaceae_RC9_gut_group* (belonging to *Bacteroides*) accounted for the largest proportion of the cecum microbial community. At these two levels, compared with the HRFI hens, the LRFI group showed a decrease in the relative abundance of *Bacteroides*, and the same trend was shown at the phylum level.

LEfSe analysis was applied to find the significant differentially abundant ASVs for the entire microbiota at levels from phylum to genus (*P < 0.05*; LDA > 3.0). As shown in Fig. 6D, *Clostridia, Oscillospirales, Ruminococcaceae, Butyricicoccaceae, Rikenellaceae_RC9_gut_group and Actinobacteria* were selected as biomarkers with substantial differences between the two groups. Compared with the HRFI group, the relative abundances of *Clostridia, Oscillospirales, Ruminococcaceae, and Butyricicoccaceae* were increased significantly in the LRFI groups, while the HRFI groups showed an increased abundance of *Rikenellaceae_RC9_gut_group* and *Actinobacteria*. In summary, LEfSe in the cecum showed that the LRFI group had a decreased relative abundance of pathogenic bacteria, while the relative abundance of probiotics was increased. These results indicated that LRFI hens had many dominant microflora in their intestines and the potential to reverse the disorder of intestinal bacteria caused by high feed intake.

### 3.7 Untargeted metabolomics analysis in the liver

LC-MS was used to detect the differences and small molecule metabolites in the liver between the LRFI hens and the HRFI hens. The total number of metabolites expressed in liver samples of different RFI laying hens was 645, which the total number of metabolites was 637, the number of DEMs in the HRFI group was 5. The number of DEMS in LRFI laying hens was 3 (Fig. 7A). PLS-DA and OPLS-DA were performed for each group of data after pretreatment. Student’s t test was used for all analyses and the confidence level was >0.95. PLS-DA, OPLS-DA scores and cluster analysis of DEMs were significantly different between the LRFI group and between the HRFI group (Fig. 7C-E). These results suggest that there were significant differences in the levels of metabolites in two groups.

**Fig. 7.**
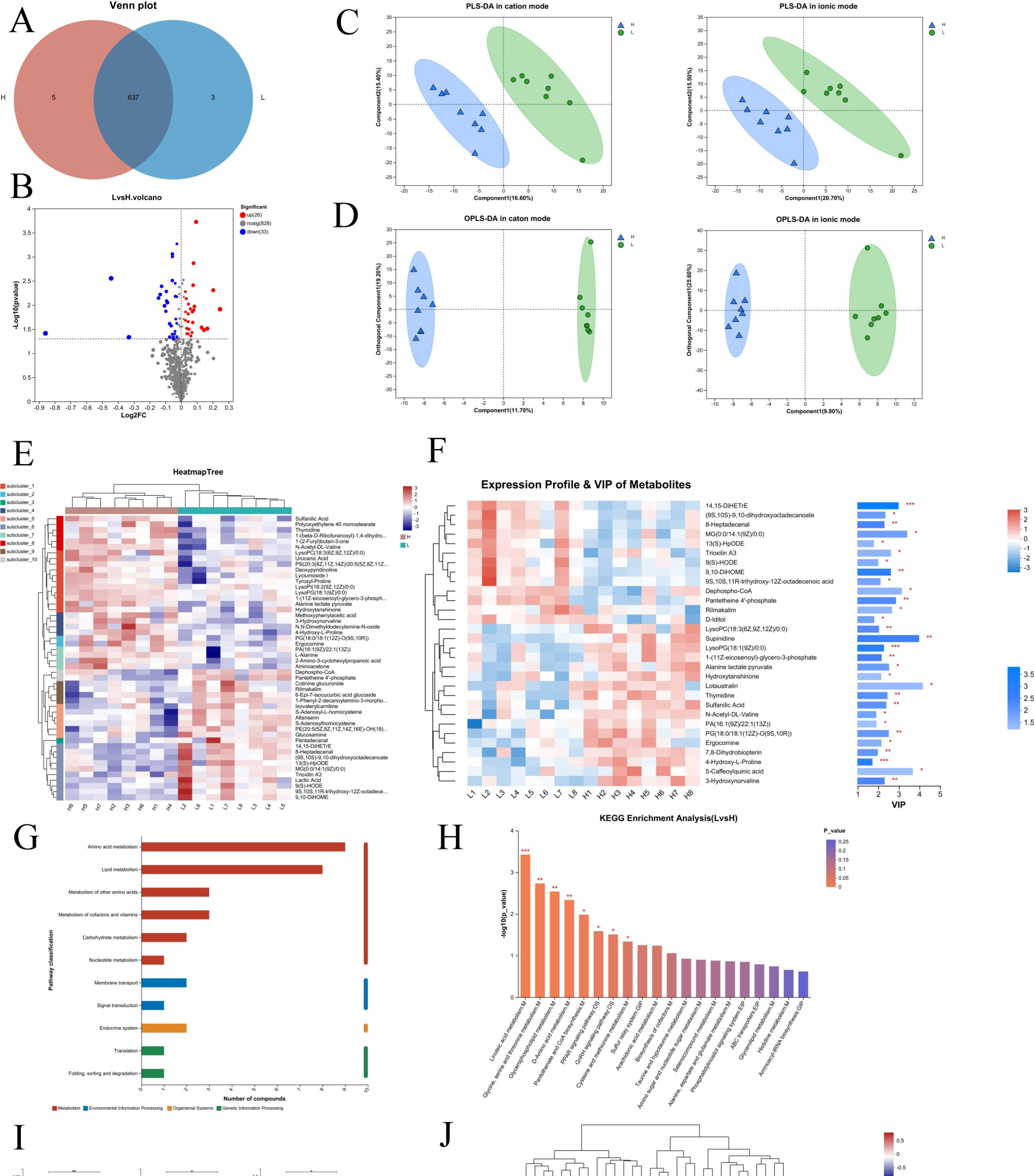
Untargeted metabolomics analysis of liver in the LRFI and HRFI groups (n = 8 per group). (A) Venn diagram of DEMS; (B) Volcano plot (combination of positive ion and negative ion); (C) PLS-DA analysis (combination of positive ion and negative ion); (D) OPLS-DA (combination of positive ion and negative ion); (E) DEMs expression in cluster heatmap; (F) Analysis of VIP values in metabolites (combination of positive ion and negative ion); (G) KEGG annotation analysis (combination of positive ion and negative ion); (H) KEGG pathway enrichment analysis (combination of positive ion and negative ion). (I) Violin Plot representation of key DEGs. (J) Correlation analysis of cecum microbes and liver metabolites. (\**P < 0.05, **P < 0.01, ***P< 0.001*).

The variable importance in projection (VIP) score map based on the OPLS-DA model could intuitively visualize the importance of DEMs. The DEMs with VIP scores >1 and *P < 0.05* were selected as metabolites with high influence on the classification model for subsequent analysis. The results showed that the top 30 metabolites with significant differences were identified between different groups (Fig. 7F). In contrast to the HRFI group, the VIP values exceeding 1 for 10 metabolites in the LRFI groups were listed as 14,15-DiHETrE, LysoPG(18:1(9Z)/0:0), 4-Hydroxy-L-Proline, 9,10-DiHOME, Supinidine, 1-(11Z-eicosenoyl)-glycero-3-phosphate, 8-Heptadecenal, 3-Hydroxynorvaline, Pantetheine 4’-phosphate, LysoPC(18:3(6Z,9Z,12Z)/0:0).

A volcano map was constructed to visualize the contribution of potential biomarkers between two groups (Fig. 7B). Each point in the picture represents a metabolite. Based on the abscissa is log _2_(fold change) and −log _10_ *P-*value, a total of 59 known DEMS were identified in liver samples of different RFI laying hens in both cationic and anionic modes.

Functional annotation analysis was performed on DEMs found in positive and negative ion modes by KEGG. The DEMs annotated in liver samples of both RFI laying hens were mainly involved in metabolism, including amino acid metabolism, lipid metabolism, metabolism of cofactors and vitamins and carbohydrate metabolism (Fig.7G). They also take part in environmental information processing, such as membrane transport and signal transduction, as well as organismal systems, specifically the endocrine system. In addition, they were associated with genetic information processing, including translation and folding, sorting and degradation. Among these functional pathways, the most enriched pathways were “Metabolism,” particularly amino acid metabolism and lipid metabolism, which are the focus of our later investigations due to their significant enrichment in metabolites (Supplemental information).

KEGG pathway enrichment analysis was conducted for the selected metabolites and the biological pathways in which the DEGs were involved. As shown in Fig. 7H, the results showed that the DEGs were mainly significantly enriched in biosynthesis of metabolism, specifically linoleic acid metabolism, glycine, serine and threonine metabolism, glycerophospholipid metabolism, D-amino acid metabolism, pantothenate and CoA biosynthesis, and cysteine and methionine metabolism. Other significantly enriched pathways included the PPAR signaling pathway and GnRH signaling pathway (*P<0.05*). Among them, L-alanine, LysoPI(18:2(9Z,12Z)/0:0) and PS(20:3(8Z,11Z,14Z)/20:5(5Z,8Z,11Z,14Z,17Z)) were found to align with the significantly enriched main KEGG pathways mentioned above. These metabolites are considered key DEGs in this experiment (Fig. 7I).

### 3.8 Correlation of the microbiota and metabolites

To investigate the potential functional connection between intestinal bacteria and liver metabolism, a correlation heatmap was set up using Pearson’s correlation analysis (Fig. 7J). The data showed that the relative abundances of *Odoribacter*, *Bacteroides* and *Ligilactobacillus* were positively correlated with N,N-dimethyldodecylamine-N-oxide; *Bacteroidales* and *Megamonas* were positively correlated with 1-(beta-D-ribofuranosyl)-1,4-dihydronicotinamide. However, the relative abundances of *Ruminococcus_torques* and *Shuttleworthia* were negatively correlated with cotinine glucuronide and 1-phenyl-2-decanoylamino-3-morpholino-1-propanol. It is worth noting that the relative abundance of the beneficial bacterium *Butyricicoccus* was significantly higher in the LRFI group than in the HRFI group (Fig. 6D). There was a significant negative correlation between the expression level of LysoPI (18:2(9Z,12Z)/0:0) in the liver and the abundance of *Butyricicoccus*. Additionally, the expression level of LysoPI (18:2(9Z,12Z)/0:0) in the liver of the LRFI group was also significantly lower than that of the HRFI group.

## 4. Discussion

Due to feed costs account for 60-70% of the total farming expenses, feed efficiency has always been one of the crucial factors determining the economic viability of poultry production. Studies have found that as the laying period progresses, particularly in the late stages of egg laying, there is a noticeable decline in feed utilization efficiency due to the physiological aging of hens^[32]^. Therefore, elucidating the genetic mechanisms of late stage egg production in hens and extending the laying period are important breeding objectives. Currently, there has been extensive research on RFI, but research on RFI in poultry remains limited^[33]^. The feeding behavior of laying hens is a complex neuroregulatory network, and studying the underlying mechanisms of biological systems cannot be fully understood through single omics approaches, such as individual genes or proteins. Therefore, in this experiment, a multiomics approach was employed to comprehensively assess gene expression, metabolic profiles, and microbial community characteristics within the organism. This approach is better suited to elucidate the molecular level regulatory mechanisms of RFI in hens during the late stage of egg production and supplies a theoretical basis for selecting and breeding RFI traits in laying hens.

Research has shown that selectively breeding for RFI essentially involves reducing feed intake while maintaining body weight gain to improve feed efficiency^[34, 35]^. In this study, we performed biostatistical analysis on the production performance and egg-laying performance of all chicken flocks in poultry farm. We ultimately identified 1192 hens for correlation analysis between RFI and other indicators. The results revealed a highly significant positive correlation between RFI_30-31_ and FI_30-31_, while no significant correlations were found with body weight traits or egg production traits. This finding is consistent with previous studies conducted on laying hens^[31, 36]^, broilers^[16, 37]^, and ducks^[38]^. In summary, we confirm that RFI is independent of growth traits and can effectively reduce feed intake and improve feed efficiency without affecting body weight and egg production. In this experiment, while ensuring similar egg production rates between the high and low RFI groups, we further measured RFI in hens at 67-69 weeks of age. We then classified the experimental population based on individual RFI values during this period into high (RFI > Mean + 0.5 SD) and low (RFI < Mean −0.5 SD) RFI groups, each consisting of 10 hens. These groups will be used for later biological analyses.

The quality of eggs plays a crucial role for both producers and consumers during storage, processing, and consumption processes. Consistent with the findings of Sun^[39]^, our study also found no significant differences between the high and low RFI groups in terms of ESI, ESC, and EW. Therefore, selective breeding for RFI during the late stage of egg production is possible, as it can reduce feed consumption without significantly affecting egg quality.

RFI reflects the difference between the amount of feed consumed by hens and their expected feed intake. Body size traits such as body weight, body length, chest circumference and chest width are closely related to the growth rate and size of chickens. By studying the relationship between body size traits and RFI in laying hens, we can further understand the differences in feed efficiency and improve production economic performance. Currently, research has shown a negative correlation between body size traits and RFI in poultry^[40]^, indicating that larger poultry can utilize feed more efficiently under the same feed consumption. Our research findings revealed a significant difference in chest width between the two groups of hens. Compared to the HRFI group, the LRFI group showed a wider chest width. This is also consistent with the findings of Wen et al.^[41]^, where a decrease in RFI values was associated with an increase in chest width in broilers.

The digestive tract of poultry mainly consists of the crop, proventriculus, gizzard, liver, intestine^[42]^. The liver’s digestive function involves utilizing bile synthesized by the body to activate pancreatic lipase, facilitating the digestion and absorption of dietary starch in the intestine^[43]^. Research has found that the length of digestive organs is typically correlated with the feed intake capacity and feed conversion efficiency of poultry. Longer lengths of digestive organs indicate a larger surface area, providing more absorption areas and enhancing the digestion and nutrient absorption capacity of feeds. This ultimately leads to improved feed utilization efficiency^[44]^. The weight of digestive organs reflects the level of development and health of the poultry digestive system. Heavier digestive organs may indicate greater digestive capacity and feed utilization efficiency^[45]^. In our results, we observed that the length and weight of the digestive organs in the LRFI group were higher than those in the HRFI group. These findings indicate that evaluating the length and weight of the digestive organs in laying hens can provide valuable insights into their feed digestion capacity, feed efficiency, and overall health status.

In the current study, blood parameter concentrations associated with feed intake, growth, nutrient allocation, and utilization served as potential physiological indicators of feed efficiency^[46]^. ALT and AST are primarily found within liver cells, with minimal amounts in the serum. When the liver is damaged, the normal structure of liver cells is disrupted, leading to a significant increase in ALT and AST levels in the blood. In poultry, elevated levels of LDL-C indicate disturbances in lipid metabolism, which may contribute to the occurrence of metabolic-related diseases^[47]^; HDL-C, as it aids in the removal of LDL-C from the body and transports it to the liver for metabolism, is generally considered “good cholesterol”^[48]^. In this study, we observed that the AST concentration was significantly lower in the LRFI group than in the HRFI group, while the HDL-C content was significantly higher in the LRFI group. This suggests that the lipid metabolism of laying hens in the LRFI group tends to be healthier. Additionally, the results of H&E staining and Oil Red O staining indicated that liver cells and nuclei in the HRFI group experienced some degree of damage, with the accumulation of lipid droplets in the liver. The presence of fat accumulation in the liver may signify the existence of fatty liver, which can affect feed intake in laying hens and reduce feed utilization efficiency to some extent.

ACTH, through stimulating the production of glucocorticoids, participates in regulating stress responses and metabolic processes. Previous findings^[49, 50]^ in poultry suggested that the ACTH levels in LRFI chickens were significantly lower than those in HRFI chickens, which is consistent with our research results. LEP is a hormone secreted by adipose tissue that reduces food intake by stimulating the satiety center. Research has shown that animals with lower RFI have higher levels of circulating hormones and their associated gene expression^[51]^. INS is the only hormone in the body that lowers blood glucose levels and promotes glycogen, fat and protein synthesis. Studies have found that the LRFI group has higher concentrations of INS, and this may be due to the relationship between INS expression and visceral fat deposition^[50]^. However, Karisa et al. found a tendency toward higher INS concentrations in HRFI individuals^[52]^. Although the concentrations of LEP and INS did not reach a significant difference level between the high and low RFI groups, our results indicated a higher trend in the LRFI group. Metzler^[53]^ through their study confirmed that there is a correlation between serum metabolite changes and RFI variation in chickens, and serum metabolite variation can be used to predict changes in RFI. Therefore, based on our study, AST, HDL-C, and ACTH in chicken serum can serve as candidate indicators for late phase RFI in laying hens. It is important to note that RFI is influenced by various factors, including hormones, appetite regulating neural pathways, dietary components, and environmental factors. Therefore, further research and investigation are needed to understand the specific relationship between serum biochemical indicators and RFI.

The liver is the largest parenchymal organ in chickens. By analyzing the transcriptome of livers, we can gain a deeper understanding of the dynamic processes of gene expression and reveal gene regulatory networks related to metabolism, immunity, and growth^[54]^. Xu^[55]^ conducted transcriptome sequencing of liver tissues from chickens with different feed efficiency and found that the DEGs in the liver were mainly associated with appetite, cellular activity, and lipid metabolism. In this study, high throughput RNA-seq technology was used to obtain transcriptomic data from the liver tissues of high and low hens (70 weeks of age). The alignment rate in previous transcriptomic studies on chickens ranged from 64.00% to 85.00%^[56]^, while the sequencing results of this study revealed an average of forty million reads or more per individual, with an average alignment rate of over 90%. These high quality reads and alignment rates ensure the accuracy and reliability of subsequent differential gene expression analysis. In total, 268 DEGs (167 upregulated genes and 101 downregulated genes) were identified from the sequencing data in this experiment. To validate the functional roles of these DEGs, KEGG pathway analysis was performed on the 268 DEGs, and the top 20 enriched results are presented. The results indicated that the main enriched pathways of DEGs were as follows: thyroid hormone synthesis, pancreatic secretion, mineral absorption, dorsoventral axis formation, cholesterol metabolism, bile secretion and the PPAR signaling pathway. These pathways mainly involve the endocrine system, digestive metabolism, cellular development, and immune response. Similar to previous studies^[55, 57]^, our results suggest that the enriched DEGs primarily regulate the RFI phenotype by affecting hormone secretion, regulation and digestive metabolism. In other words, these DEGs associated with these processes may serve as potential functional genes influencing RFI. In this study, we found that *ADCY2, ADCY8, and CCKAR* were significantly upregulated in the liver of LRFI laying hens, while *ACSS2, FABP1,* and *FABP4* were significantly downregulated. *ADCY2* and *ADCY8* are transmembrane enzymes involved in intracellular signal transduction and regulate energy metabolism. Current research has found that specific variants of the *ADCY* gene family are associated with suppressive feeding behaviors. These variants may affect the activity of adenylate cyclase 2, thereby altering intracellular cAMP levels and subsequently influencing the regulation of appetite and feeding behavior. Moreover, *ADCY* genes are also thought to be involved in the regulation of lipid synthesis, playing a role in inhibiting fat synthesis or increasing fat breakdown^[58]^. *ACSS2* catalyzes the generation of Acetyl-CoA, providing substrates for energy production and the synthesis of fatty acids^[59]^. Therefore, the high expression of *ACSS2* may promote fatty acid synthesis and energy supply, thereby enhancing the feed intake and fat synthesis of livestock and poultry. *CCKAR* is a member of the cholecystokinin (*CCK*) receptor family, which is associated with appetite regulation and feeding behavior^[60]^. According to existing research, *CCKAR* is believed to be associated with suppressive feeding behavior. *FABP1* and *FABP4* are mainly expressed in the liver and can promote the uptake, transport, and oxidative metabolism of lipids in hepatocytes. Studies have shown that high expression of the *FABP* gene family may result in increased lipid accumulation in hepatocytes, leading to the release of signaling molecules or stimulating factors that affect appetite regulation centers, thus increasing feed intake and fat deposition in the body^[61]^. Similar to previous studies on key DEGs related to RFI in livestock and poultry^[59, 60, 62]^, we hypothesize that *ADCY2, ADCY8, CCKAR, ACSS2, FABP1,* and *FABP4* are key candidate genes influencing RFI in late-stage egg laying hens. To validate the accuracy of the predicted DEGs in this study, we conducted qRT‒PCR on these six candidate genes. We found a high degree of consistency between the RNA-Seq based predictions and the experimental validation results, indicating the high accuracy of our study findings. Although this study identified 30 potential functional genes related to RFI enriched in pathways, the biological replication used in this study was limited. Further investigations on RFI should involve larger populations and employ multialgorithm approaches to elucidate its genetic mechanisms.

The liver is a central organ for the metabolism of various tissues. Through functions such as bile secretion, protein metabolism, carbohydrate metabolism and lipid metabolism, it helps the body digest, absorb nutrients and maintain metabolic balance^[63]^. Based on the DEMs, the analysis of potential metabolic pathways was measured, which was helpful in revealing the mechanism of different RFI groups. The results showed that the metabolites were mainly enriched in linoleic acid metabolism, glycine, serine and threonine metabolism, glycerophospholipid metabolism, D-amino acid metabolism, pantothenate and CoA biosynthesis, pantothenate and CoA biosynthesis, the PPAR signaling pathway, the GnRH signaling pathway and cysteine and methionine metabolism. These pathways are mainly involved in lipid metabolism, amino acid metabolism, metabolism of cofactors and vitamins, and the endocrine system. They primarily participate in the body’s lipid metabolism and nutrient intake, collectively maintaining normal physiological functions and homeostasis in the body. Among the enriched metabolites, lipid metabolism, amino acid metabolism and carbohydrate metabolism are closely related to poultry digestion, utilization and feed intake which have been proven to be closely associated with RFI^[64, 65]^. Combining our previous findings from liver histological staining and transcriptome analysis, we have also preliminarily observed, when egg production rates are similar, hens in the high feed intake group tend to have higher lipid deposition in the late stage of egg production compared to the low feed intake group. In this study, our DEMs were mainly enriched in the KEGG pathways of lipid metabolism and amino acid metabolism. For example, L-alanine, enriched in glycine, serine and threonine metabolism, has been proven to be involved in protein synthesis and energy metabolism in the body. Additionally, L-proline can act as a flavor enhancer, increasing chicken interest and appetite for food, thereby promoting increased feed intake^[66]^. LysoPI(18:2(9Z,12Z)/0:0) and PS(20:3(8Z,11Z,14Z)/20:5(5Z,8Z,11Z,14Z,17Z)), which are enriched in glycerophospholipid metabolism, are phospholipids that primarily participate in cellular membrane composition and signal transduction and serve as important substrates for fatty acid synthesis. Liu^[67]^ found that PS[18:2(9Z,12Z)/20:4(5Z,8Z,11Z,14Z)] and PS[18:0/20:4(8Z,11Z,14Z,17Z)] may indicate that the LRFI group has enhanced cell membrane integrity, fatty acid transport, nutrient absorption, immune function and signal transduction to improve feed efficiency. In this study, L-alanine, LysoPI(18:2(9Z,12Z)/0:0) and PS(20:3(8Z,11Z,14Z)/20:5(5Z,8Z,11Z,14Z,17Z)) were significantly negatively correlated with LRFI groups, indicating that changes in metabolites in the liver resulted in a difference in feed utilization. However, there is still limited research on the relationship between amino acids and phospholipids with RFI, and further in-depth studies are warranted.

Numerous studies have demonstrated a close association between the species and abundance of cecal microbiota and feed utilization efficiency in poultry^[68, 69]^. *Firmicutes* and *Bacteroidetes* are the two major phyla in the gut microbiota. Previous studies have indicated that poultry with different feed conversion efficiency exhibit variations in the ratio of *Firmicutes* to *Bacteroidetes*^[70]^. In poultry, with low feed utilization efficiency indicated a decrease in the abundance of *Bacteroidetes* and an increase in the abundance of *Firmicutes* in the intestinal microbiota. Due to the decreased ratio of *Firmicutes* to *Bacteroidetes*, which inhibits the absorption and utilization of energy in poultry, leading to lipid accumulation or the occurrence of fatty liver^[71, 72]^. In this study, we found that the bacterial community structures in caecum with different RFI phenotypes were also significantly different. Compared to the HRFI group, the LRFI group showed an increase microbiota diversity and an elevated ratio of *Firmicutes* to *Bacteroidetes*. Through LEfSe analysis, we found some representative species as biomarkers to distinguish the gut microbial community between the two groups. In the LRFI group, the intestinal flora was mainly enriched in *Clostridia*, *Oscillospirales*, *Ruminococcaceae*, and *Butyricicoccaceae*, which is also consistent with previous studies in relation to RFI^[73, 74]^. Studies have found that most of the microbiota associated with feed conversion efficiency are involved in cellulose decomposition, fermentation, and metabolism^[75]^. *Clostridia*, *Ruminococcaceae* and *Butyricicoccaceae* are recognized as beneficial bacteria can breakdown cellulose ^[76]^ and produce short chain fatty acids (SCFAs: mainly include propionic acid, acetic acid, and butyric acid) ^[77]^. These fatty acids can be absorbed by the intestinal tract of laying hens, providing energy and participating in bile acid metabolism. SCFAs can further influence the absorption of fat and cholesterol by modulating bile acid levels^[78]^, maintaining the balance of the gut microbiota, and positively affecting digestion and nutrient absorption, thereby reducing feed intake and improving feed utilization efficiency^[79]^. Additionally, acetic acid and butyric acid have been shown to inhibit animal feed intake and reduce fat deposition^[80, 81]^. Therefore, we can tentatively speculate that the effects of reducing lipid accumulation and improve feed utilization efficiency in laying hens are attributed to the presence of SCFAs in the intestinal tract. However, the complexity and diversity of the microbial community make it challenging to accurately determine causal relationships between specific bacterial groups and physiological indicators. Further research is needed to explore and validate these findings.

The gut-liver axis indicates a tight relationship between intestinal bacteria and the liver^[82, 83]^. By incorporating the gut-liver axis theory, a correlation analysis was conducted between the major gut bacteria and DEMs in the liver. The related analysis was set up to clarify the association between cecum bacteria and liver metabolites between the LRFI and HRFI groups. Notably, the results showed that *Butyricicoccus* was negatively correlated with LysoPI(18:2(9Z,12Z)/0:0). In addition, LysoPI(18:2(9Z,12Z)/0:0) played an active role in fat synthesis, while the production of SCFAs was significantly positively correlated with *Butyricicoccus*^[84]^. These results are consistent with the conclusions of the analysis of intestinal microbial species diversity, indicating that the difference in gut microbiota and hepatic metabolites can affect the lipid metabolism and feed intake of laying hens.

## 5. Conclusion

The RFI is independent of growth traits, and selecting individuals with low RFI can improve feed efficiency without affecting the laying performance and egg quality of hens. In this study, it was found that under similar egg production rates, hens in the HRFI group had lower nutrient digestion capacity, disrupted lipid metabolism, and lower feed utilization efficiency than those in the LRFI group. Further research and validation are needed to investigate the effects of AST, HDL-C, and ACTH on feed efficiency in the late laying period through serum biochemical index analysis. Through multiomics analysis, genes such as *ADCY2, ADCY8, CCKAR, ACSS2, FABP1,* and *FABP4* and metabolites such as LysoPI(18:2(9Z,12Z)/0:0) were identified as key genes and biomarkers related to RFI in laying hens during the late laying period. These findings are important for developing strategies to improve feed efficiency in Rhode Island Red pure-line chickens during the late laying period. Additionally, in-depth investigations such as histological observations, enzyme activity analysis, and integration of multiomics techniques are needed to unravel the physiological and biochemical mechanisms of digestive organs that impact feed utilization efficiency in poultry.

## CRediT authorship contribution statement

Zhouyang Gao: Carried out the experiments, Data curation, Writing. Chuanwei Zheng: Supervision. Zhiqiong Mao and Jiangxia Zheng: Revision, suggestion. Dan Liu and Guiyun Xu: Project administration.

## Declaration of competing interest

The authors have declared that no conflict of interest.

## Acknowledgements

This study was supported by the National Key Research and Development Program of China (2022YFD1300100).

## Notes

### Competing Interest Statement

The authors have declared no competing interest.

